# Role of RBC membrane protein palmitoylation in regulation of molecular topology and susceptibility to *Plasmodium falciparum* invasion

**DOI:** 10.1101/2020.01.21.913855

**Authors:** Soumya Pati, Preeti Yadav, Geeta Kumari, Rex D.A.B, Sangam Goswami, Swati Garg, T.S. Keshava Prasad, Sivaprakash Ramalingam, Shailja Singh

**Author notes:** Corresponding authors Correspondence: Dr. Shailja Singh, Associate Professor, Special Centre for Molecular Medicine, Jawaharlal Nehru University, New Delhi-110067, India. E.mail; Ph No: 011-2670 4559 Ext-3038. Joint first authors (Soumya Pati, Preeti Yadav, Geeta Kumari).

## Abstract

Squeezability of biconcave RBC raises a fundamental query, about, how it can restructure its bendable cytoskeleton for efficient micro-circulation. We report for the first time, the existence of dynamic palmitoylome in RBC composed of 118 palmitoylated proteins that reduced to 42 upon treatment with 2BP, a generic inhibitor of palmitoylation. In-depth analysis revealed that Semaphorin7A, CR1 and ABCB6, the known RBC receptors for *P. falciparum* were reduced to negligible in 2BP-treated RBCs, suggesting palmitoylation-dependent recruitment of parasite-specific receptors. Interestingly, Kell, a single disulphide-linked co-partner in Kell-Kx complex was undetected in 2BP-treated RBCs, while Kx remained intact. RBCs-blocked with anti-Kell antibody demonstrated signficant reduction in parasite invasion, thus suggesting it as a receptor proto-type for *P. falciparum* invasion. Finally, reduced expression of Kell in palmitoylated protein pool of sickle-cell RBC ghost, with its diminished surface representation in these RBCs, proposed Kell, as one of the novel receptor-prototype for *P. falciparum* invasion.

## Introduction

It’s really intriguing that RBC represents incredible squeezability with super-bendable cytoskeleton, as evident by its efficient microcirculation in tiny capillaries of organs and brain to deliver oxygen^1^. Interestingly, majority of its flexibility comes from its cytoskeletal mesh, which is apparently composed of rod-shaped spectrins with size of 80 nm, connected at junctions of other structural protein complexes, including F-actin, adducin, protein4.1, tropomodulin1 *etc*^2^. Known for its elastic activity as a string, spectrins help RBCs to manage sheer mechanical stress, as with 8-μm diameter of discoid shape they maintain high-speed circulation through capillaries of 2 μm^3^. Earlier, cell mechanics-based studies highlighted that, stretchability of spectrin tetramer has been found to be raised up to 2.5 folds with the bilayer tension of 500 pN/*μ*m under hydrodynamic pressure that is assumed to be good enough to open the mechanosensitive channels with no damage to lipid bilayer^4^. We hypothesized that such extraordinary deformability in RBC might be due to the S-palmitoylation process, a lipid-based PTM at its membrane that actively remodels its cytoskeleton and assigns unconstrained shape determinability to the same. Briefly, dynamicity of S-palmitoylation involves post-translational incorporation of palmitic acid to conserved cysteines in membrane-embedded proteins^5, 6^. This is achieved via easy accessibility of aspartate–histidine– histidine–cysteine (DHHC) signature motif containing, membrane-bound palmitoyl acyl-transferases (PATs) to their protein substrates^7, 8^. Following palmitoylation, several crucial events occur at membrane, including; i) *spatiotemporal organization of proteins*, ii) *increased cross-talk of transmembrane proteins to their co-partners*, iii) *conformational modulation leading to formation of macro-molecular platforms*, and iv) *finally prevention of ubiquitination at lysine residues proximal to cysteine-rich motifs of palmitoylation sites*^6^. Though, scanty evidences suggested that, few RBC structural proteins namely, Protein4.1, Protein4.2, Band3 and Ankyrin are found to be palmitoylated^9, 10^, however there is paucity in the knowledge about the palmitoylation-based structural remodeling and tethering of surface proteins in RBC membrane. Plausibly, skeletal proteins, such as Band3, ABCB6, CD55, Basigin, DARC, Semaphorin7A and CR1 have been known to act as host-derived receptors for *P. falciparum* interaction and invasion^11, 12^. Thus, to bridge the gap in the understanding, we have addressed the unsolved queries, such as; i) whether there is any existence of dynamic palmitoylation process in RBC membrane, ii) how it imparts remodelling of membrane proteins and iii) its impact on assignment of docking sites for parasite interaction and invasion. Towards these objectives, we have deciphered for the first time RBC-specific global palmitoylome, using chemico-proteomics tools, identified the differential palmitoylome using LC-MS/MS and annotated their functional significance. In-depth analysis of RBC global palmitoylome depicted dynamic palmitoylation of earlier known parasite specific receptors, such as, Semaphorin7A, CR1 and ABCB6 (Langereis blood group). Further, blocking of palmitoylation in RBCs imposed substantial decrease in parasite invasion, suggesting aborted palmitoylation-mediated deregulated expression of parasite receptors on host membrane. Remarkably, we also found a novel blood group antigen Kell, a type II glycoprotein, co-expressed with its partner Kx as part of Kell-xK assembly on RBC membrane; as one of the crucial molecular determinants for *P. falciparum* invasion. Additionally, attenuated palmitoylation-dependent diminished expression of Kell on sickle-cell RBCs from SCA patients, suggested its possible role as one of the crucial host-specific receptor proto-type for *P. falciparum* invasion. Previously, Kell has also been found to be phosphorylated during *P. falciparum* invasion^13^. Concievably, this study reports for the first time dynamic palmitoylation process in RBCs that plays a well defined role in stable tethering of proteins in membrane, and governs the exposure of Kell in membrane, thereby increasing its susceptibility towards malaria parasite invasion.

## Results

### Metabolic probe-based Click reaction revealed a dynamic palmitoylation process in RBC

To identify the existence of a dynamic palmitoylation process in RBCs, click chemistry, a highly sensitive tool was performed that involves Cu(I)-catalyzed cyclo-addiction of palmitic acid alkyne (C16:0) analogues (17ODYA) to the exposed cysteines in proteome of RBC ghost^14^, which was subsequently visualized by an Azide functionalized dye (Oregon green) (Figure-1) using confocal imaging. This phenomenon was completely aborted by inhibiting the PATs by 2BP treatment, as evident by diminished fluorescence intensity suggesting stalled incorporation of labeled 17ODYA (Figure-1(a)), leading to substantial disfiguration of RBC membrane structure. Evaluation of MFI of 2BP-treated individual RBCs strongly depicted abrogated palmitoylation as compared to untreated controls (Figure-1 Right Panel). These observations suggested that RBCs harbor a dynamically active palmitoylation process at membrane that might regulate its shape and integrity.

**Figure 1.**
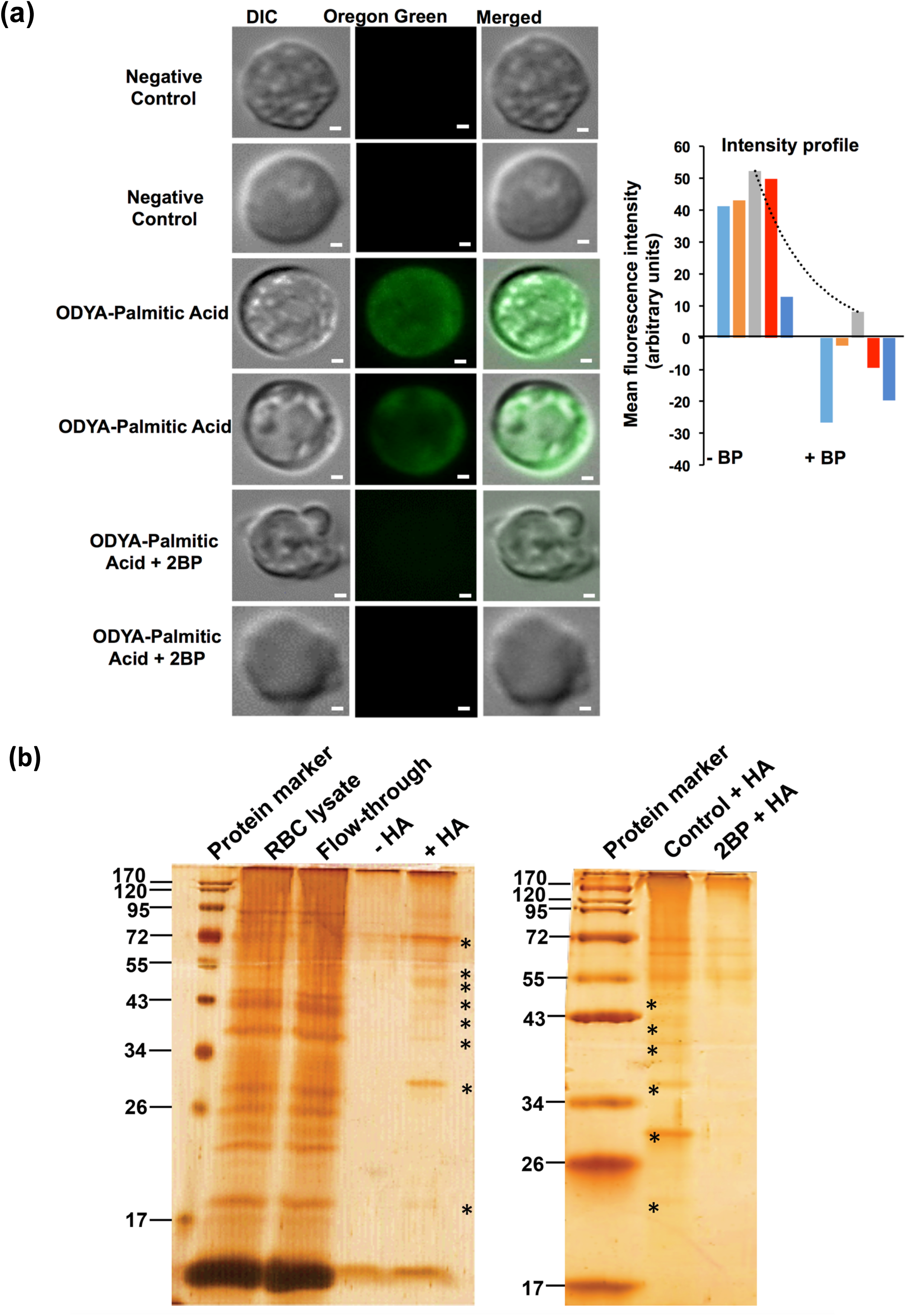
(a) Visualisation of global palmitoylome of RBC using metabolic-labelling coupled click chemistry. Scale bar: 1µM. Graph on the right shows comparative intensity profile of Oregon-green fluorescence in 5 different RBCs with and without 2BP treatment. (b) Silver-stained SDS-gel representing palmitoylated protein bands of RBC membrane lysate processed through ABE method. Bands in +HA lane corresponds to palmitoylated proteins, with-HA being the negative control. Gel on the right compares the bands of palmitoylated RBC membrane proteins with and without 2BP treatment.

### Acyl Biotin Exchange-based tool enriched global palmitoylome in RBC ghost

To enrich and quantify ghost-specific global palmitoylome, Acyl Biotin Exchange (ABE) was performed for total RBC ghost lysate and data was interpreted using silver staining of the eluted protein fractions (Figure-1(b)). Briefly, the total palmitoylated proteome was extracted from RBC ghost lysate, that followed HA treatment, exposing palmitoylated cysteine’s thiol groups that were then selectively labeled by biotin–HPDP, a thiol-reactive agent. The modified proteins were then fished out using streptavidin beads, as visualized by silver stained SDS-gels (Figure-1(b)). The results (Figure-1(b) Right Panel) represented substantial reduction in palmitoylation upon 2BP treatment in hydroxylamine treated protein pools (+HA-2BP), as majority of bands disappeared in +HA-2BP compared to +HA-control. Further, for identification and analysis of global palmitoylome in LC-MS/MS, band elutes from both 2BP treated and untreated lysates were further processed, as described in Methods Section (In Solution Digestion).

### LC-MS/MS based analysis of global palmitoylome profiling revealed dynamically palmitoylated distinct protein clusters in RBC membrane, upon treatment with 2BP

Following ABE, the eluted protein fractions were subjected to LC-MS/MS for protein identification. The result obtained was matched with CSS-Palm prediction software, as shown earlier by our laboratory^15^, which identified proteins with evident palmitoylation, containing atleast one palmitoylation site per protein in ∼90% of proteome. Based on this, we could predict 118 proteins in control sample, out of which only 42 proteins could be detected in 2BP treated RBC sample (Figure-2(a), Table-1, Table-2). For these proteins, the coverage, peptide number and score were highly reduced. To evaluate the frequency of palmitoylation in each protein, cysteine-rich palmitoylation sites predicted by CSS-Palm were catalogued, delineating a maximum number of 11 sites in a protein, namely, Phospholipid Scramblase1. Precisely, majority of proteins were found to have 1-2 palmitoylated sites (Figures-2(b)).

**Figure 2.**
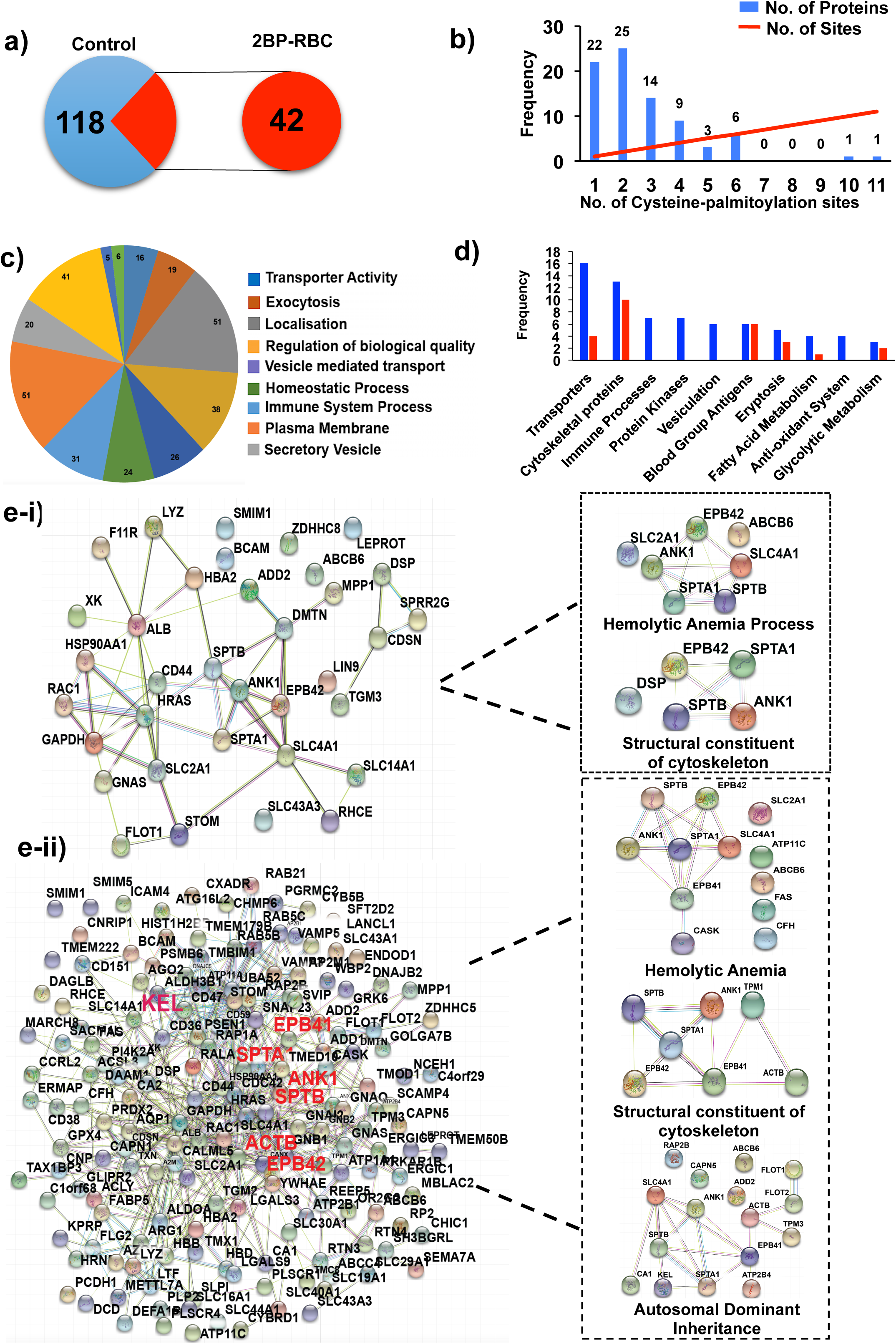
(a) Pie charts represent total palmitoylated proteins in RBC membrane, before and after 2BP-treatment respectively, determined by LC-MS/MS. (b) Bar-graph showing frequency of cysteine-containing palmitoylation sites in all predicted palmitoylated proteins of RBC ghost. (c) Venn diagram highlighting protein abundance of different functional clusters of RBC membrane palmitoylome, predicted by GO enrichment analysis. (d) Bar-graph comparing the frequency of proteins corresponding to different functional clusters in RBC samples treated with and without 2BP. (e) Protein-protein interactions were predicted between palmitoylated proteins of RBC membrane before (ii) and after 2BP (i) treatment, using STRING resource. Specific protein clusters identified in each sample are highlighted.

### Functional classification of RBC palmitoylome using GO enrichment analysis identified crucial molecular and cellular bioprocesses in RBC

The predicted palmitoylated proteins of RBC ghost were analyzed for significant GO categories (Supplementary Figure-1) and further delineated into different modules based on their functional significance (Supplementary Figure-2). GO terms including Enzyme modulators (e.g., glycolytic enzymes, kinases, ATPases, acetylcholinesterase), Transporters (e.g., Band3 that regulates Cl^-^/HCO_3-_ transport, regulating acid-base homeostasis), Transferases, Hydrolases and Transfer/carrier proteins (e.g., Calcium ATPases, Scramblases), *etc* were found to be enriched (Figure-2(c)), emphasizing their functional annotation based on palmitoylation. Figure-2(d) highlights the absence of proteins belonging to respective protein clusters, upon 2BP treatment. As evident, clusters including Immune Processes, Vesiculation, Protein Kinases and Anti-oxidant System were severely affected as they do not represent any protein upon 2BP treatment, while others including Transporters, Fatty acid metabolism, Glycolytic Metabolism, Eryptosis, and Cytoskeletal Proteins showed reduction in protein number upon 2BP treatment. Blood-group antigens do not show any reduction in protein number upon 2BP treatment but other parameters like coverage, peptide number, *etc* were affected (Table-1, Table-2).

### Palmitoylation-driven protein-protein interactions in RBC palmitoylome represented crucial macromolecular clusters

To ascertain functional protein association networks, we constructed protein-protein interaction map of both control and 2BP treated proteins and assessed the probable modules within the interactome. Remarkably, we found more number of interactions among proteins in the control group in comparison with 2BP treated ghost lysate (Figure-2(e)). The control lysate contained marked clusters of “Structural constituent of cytoskeleton”, “haemolytic anaemia” and “Autosomal dominant inheritance”, as three major clusters driven by palmitoylation, while the 2BP-treated ghost demonstrated absence of many of the structural proteins involved in cytoskeleton and haemolytic anaemia. The protein cluster representing “Autosomal dominant inheritance”, could not be detected in 2BP-treated lysate (Figure-2(e)).

### Palmitoylation-mediated membrane organization of RBC Structural proteins

To elucidate the impact of palmitoylation on skeletal and integral membrane proteins, we performed in-depth analysis of differentially palmitoylated proteins, enriched from LC-MS/MS data from 2BP-treated RBC ghost samples. It is noteworthy that, RBC cytoskeleton comprised of a unique *spectrin*-*actin* meshwork, where six triangular spectrin structures link to one actin to constitute a net-like conformation. This structural unit has three junction complexes that fix the membrane proteins like, Glycophorin C & D (GPC & GPD), Rh, Kx and DARC to the actin-spectrin lattice via Protein 4.1, and three ankyrin complexes that plug in Ankyrin to β-Spectrin at one side and Band3 and RhAG to other part of RBC surface^1^. Further, the vertical interaction of cytoskeleton to integral transmembrane proteins is facilitated by Ankyrin, 4.1 and 4.2 proteins and Band3^16, 17^. Most importantly, besides Glycophorins, all the above mentioned structural proteins, those involved in RBC structural organization were dynamically palmitoylated, as highlighted in representative box plot (Figure-3(a)) and in the scheme, representing Structural palmitoylome of RBC (Figure-3(b)). The Y-axis in the box-plot indicated protein abundance, and the data showed drastic reduction in the abundance of these proteins in ghosts treated with 2BP. Proteins such as, ANK1 (Ankyrin), EPB42 (Protein 4.2) and FLOT2 (Flotillin 2) showed upto 2-fold decrease, whereas MPP1 and Band3 showed 4-6 and 6-8 fold reduction in protein abundance respectively, upon 2BP treatment. Interestingly, among the blood group antigens, Kell, a member of type II glycoprotein, usually co-expressed with its transmembrane co-partner Kx was found to be severely affected in 2BP-treated RBC ghost as compared to untreated fraction (Figure-3(a)). Further, we have validated the impact of dynamic palmitoylation on expression of Kell in RBC ghost using immunoblotting (Figure-3(c)), with two previously reported palmitoylated RBC structural proteins, MPP1, and Protein 4.1 as positive controls (Figure-3(d)). Probing the eluted protein fractions following ABE, for Kell, showed drastic depletion of its expression in RBC ghost upon 2BP treatment, authenticating the LC-MS/MS data (Figure-3(c)). Further probing of MPP1 and Protein 4.1 in total RBC lysate, showed huge reduction in the same following 2BP treatment, validating the differential palmitoylation data obtained by LC-MS/MS (Figure-3(d)). Additionally, DHHC17 (PAT17) expression was also found to be severely depleted in 2BP-treated RBC ghost (Supplementary Figure-3(b)).

**Figure 3.**
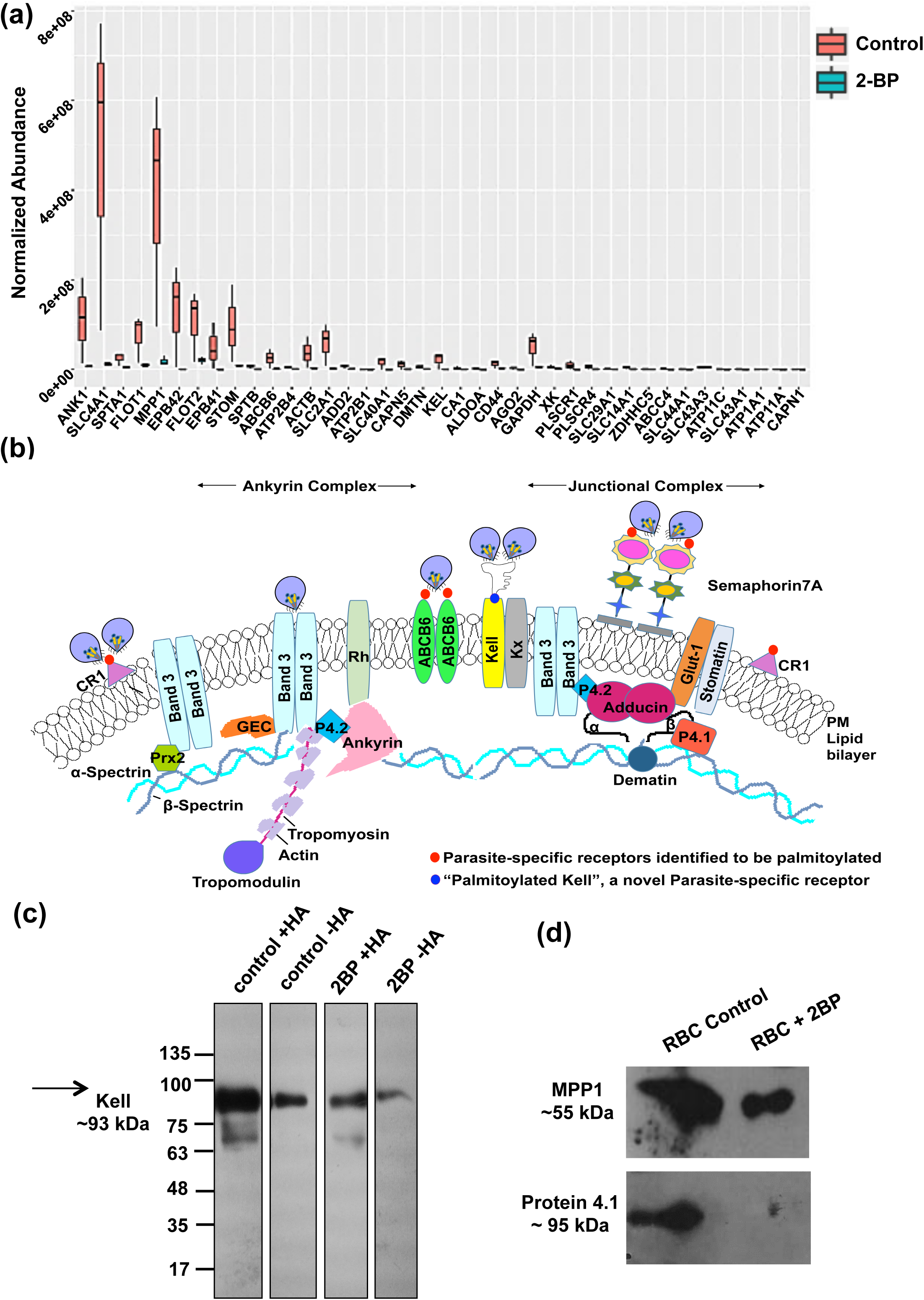
(a) Box-plot highlights the Abundance of predicted palmitoylated proteins of RBC membrane (Red box) and compares it with Protein Abundance of 2BP-treated RBC sample (green box). Vertical length of each box represents error. (b) Scheme showing structural palmitoylome identified by LC-MS/MS. (c) Immunoblot of RBC ghost palmitoylome using goat-anti-Kell IgG after ABE-based protein elution. (d) Immunoblots showing the expression of MPP1 and Protein 4.1 in RBC ghost lysate.

### Confocal imaging-based membrane localization study of Kell on RBC surface suggested its distorted tethering due to aborted palmitoylation

To evaluate the palmitoylation-dependent localized expression of Kell on RBC surface, confocal imaging was performed on both 2BP-treated and untreated unpermeabilized RBCs, using Kell-specific antibody. The results indicated disappearance of Kell on 2BP treated RBC membrane, as compared to untreated control, also evident in the MFI, which was calculated based on fluorescence intensity obtained from 50 individual RBCs (Figure-4(a)). Interestingly, there was absolutely no change detected in the localized expression of Kx, the known transmembrane co-partner of Kell ((Figure-4(b)). This finding strongly corroborated the LC-MS/MS data (Table-2).

**Figure 4.**
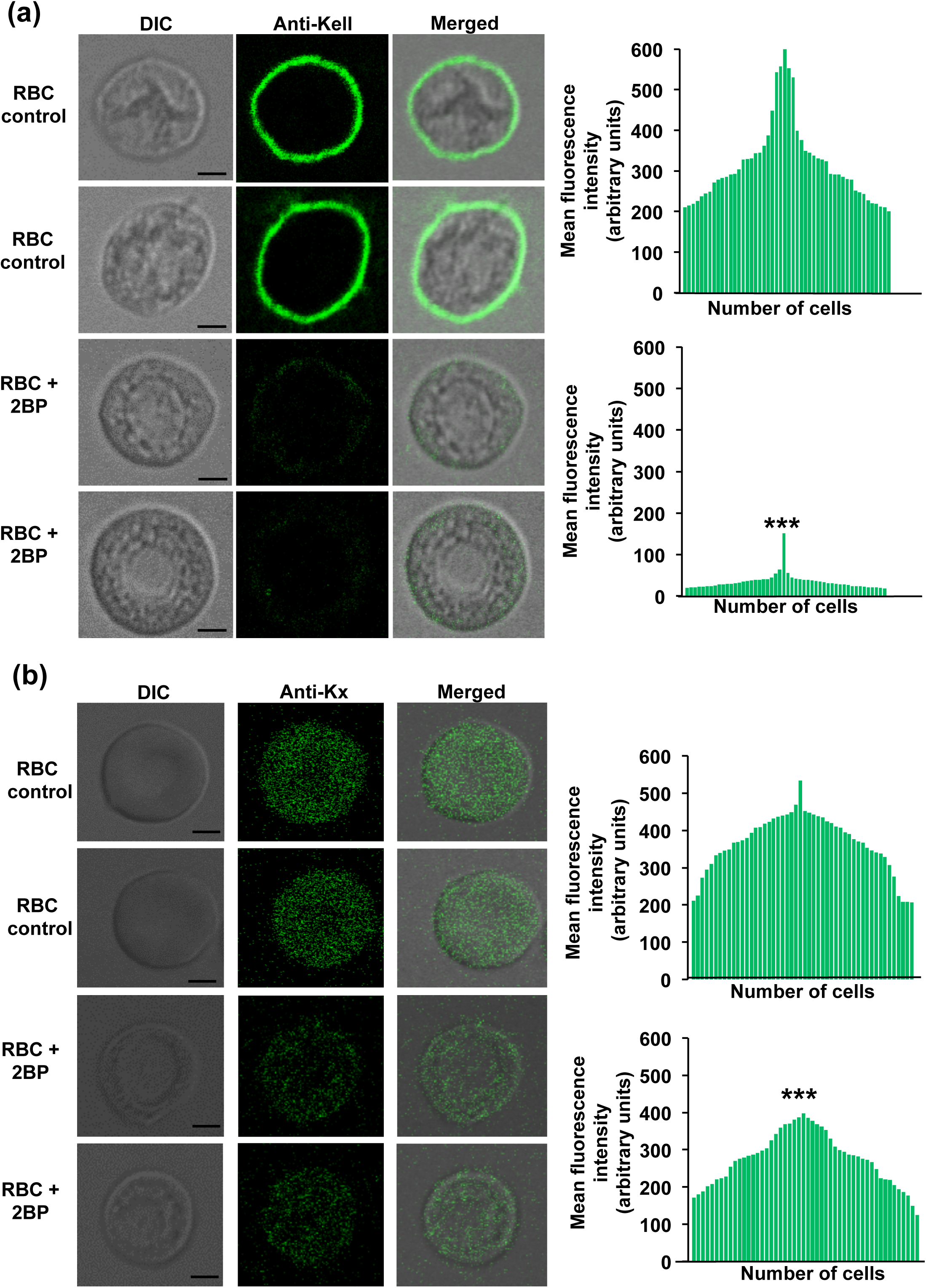
(a) Confocal images showing localisation of Kell and Kx on the surface of RBCs treated with and without 2BP. Bar-graphs represent MFI of 50 RBCs for each set. P-value <=0.0005

### Dynamic palmitoylation in RBC and palmitoylation-dependent membrane-recruitment of Kell regulate *P. falciparum* invasion

Till date, there was no information available to support any direct or indirect role of palmitoylation on parasite invasion. Thus, to evaluate the impact of dynamic palmitoylation process on parasite invasion, we treated RBCs with 2BP and performed invasion assay in the same at 1% parasitemia. The result depicted ∼70% reduction in invasion at 100µM concentration (Figure-5(a)), highlighting significant role of RBC palmitoylation in parasite invasion. Existing report suggested that Band3, the known palmitoylated structural protein in RBC has been shown to act as a host-derived receptor for *P. falciparum* invasion. Thus we asked, if dynamic palmitoylation-mediated membrane recruitment of Kell might also play the role of a receptor-prototype for parasite invasion. To understand this, RBCs were treated with anti-Kell antibody for 6hrs and were screened for number of Rings, 20-hours post-invasion in the same. The results demonstrated significant reduction in parasitemia (∼50%) (Figure-5(b)), suggesting essential role of Kell in *P. falciparum* invasion.

**Figure 5.**
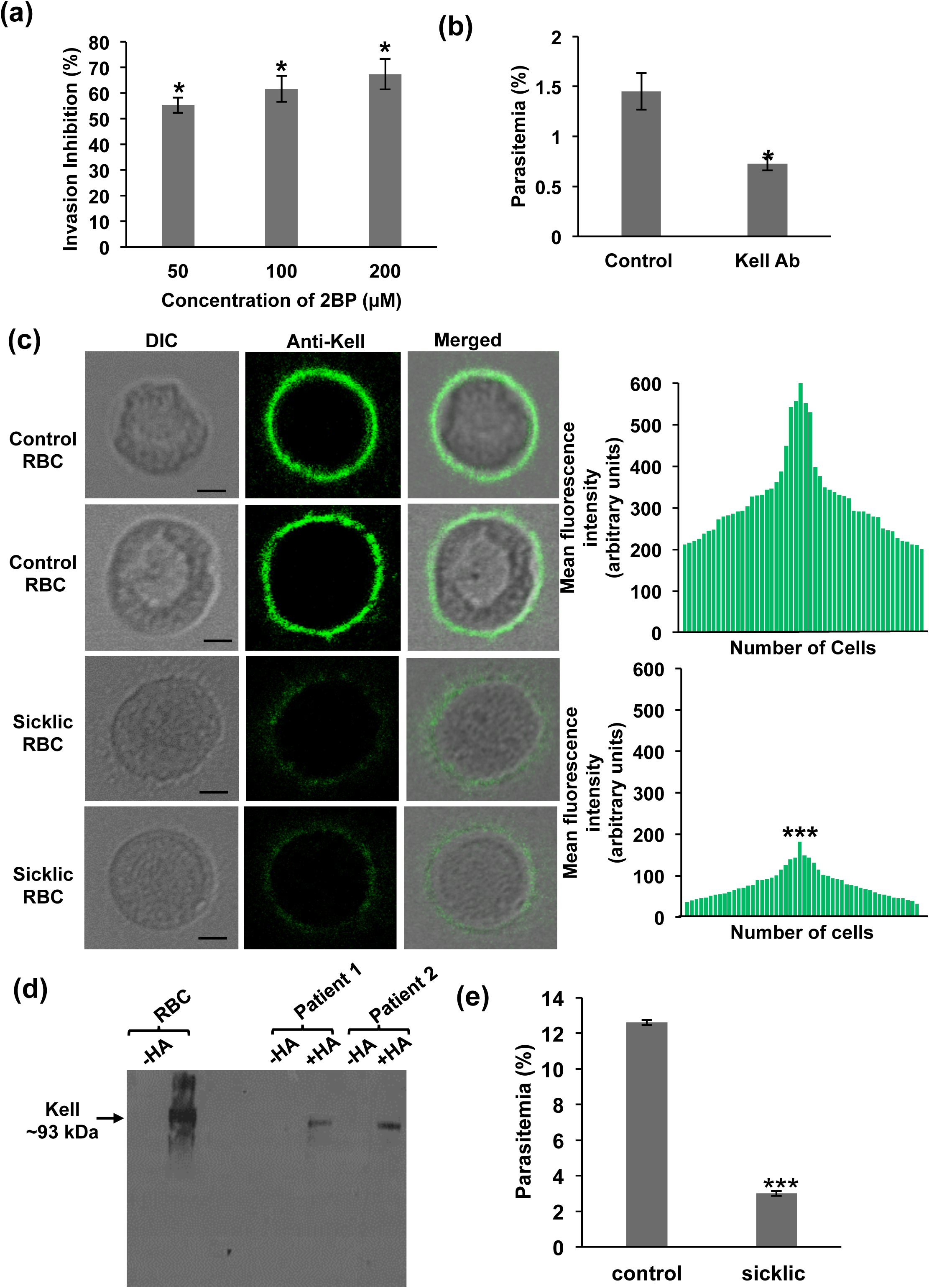
(a) Graph represent percent-inhibition of *P.falciparum* invasion in RBCs treated with different concentrations of 2BP. The data represents S.D. P-value <=0.05 (b) Graph depicts percent-parasitemia upon *P.falciparum* invasion in RBCs treated with anti-Kell IgG, taking untreated RBCs as positive control. The data represents S.D. P-value <=0.05 (c) Immunofluorescence assay comparing localisation of Kell on surface of sickle-cell RBCs in comparison to normal healthy RBCs. Bar-graphs represent MFI of 50 RBCs for each set. P-value <=0.0005 (d) Immunoblot showing the expression of Kell in sickle-cell RBC palmitoylome of 2 SCA patients with healthy RBC ghost palmitoylome as control. (e) Graph represent percent-parasitemia post-invasion of *P.falciparum* schizonts in sickle-cell RBCs, compared to healthy human RBCs. The data represents S.D. P-value <=0.0005

### Attenuated palmitoylation-mediated deregulated Kell expression in sickle-cell RBCs suggested, Kell as an essential receptor proto-type for *P. falciparum* invasion

Since expression of Kell is essential for parasite invasion, as strongly inferred by invasion results from RBCs treated with anti-Kell-antibody, we assumed that in malaria resistant RBCs, Kell expression might be severely altered due to abnormal palmitoylation. To prove this hypothesis, we then used RBC samples from 2 SCA patients, the known malaria resistant RBC model^18, 19^, and performed IFA on these patient RBCs using anti-Kell antibody. The results conferred negligible expression of Kell on sickle-cell RBCs, as shown in confocal micrographs, compared to healthy controls, also represented by MFI graph, estimated by calculating fluorescence intensity from 50 individual RBCs (Figure-5(c)). To analyze the level of palmitoylation in sickle-cell RBCs, we performed ABE to enrich the palmitoylated proteins in patient RBC ghosts, and evaluation of Kell was done using immunoblotting (Figure-5(d)). As expected, HA-treated sickle-cell RBCs demonstrated prominent reduction in Kell expression (Figure-5(d)). Further analysis of parasite invasion in sickle-cell RBCs represented severely impaired invasion phenotype, supporting earlier studies. To highlight, sickle-cell RBCs did not show any change in the expression of Kx, the glycoprotein partner of Kell (Supplementary Figure-4). This study suggested that Kell is an essential host-derived factor for *P. falciparum* invasion.

## Discussion

RBC possesses an extraordinarily fluid membrane with inherent endurance towards huge mechanical stress encountered during its microcirculation that can cover a distance upto 500km. With a diameter of 8 μm and thickness of 2–2.5 μm at periphery, it squeezes through microcapillaries of <3 μm diameter and splenic sieves of 1 μm openings^20, 21^. Such degree of flexibility by RBC cytoskeleton raises a very fundamental question, such as; how RBC being an inert cell could afford to perform structural modeling of proteins to maintain its membrane fluidity along with rheological attributes. It is noteworthy that, thioesterification of C-16 fatty acyl chains at both conserved and exposed cysteine thiols, enforces tilting, tethering and protein-protein interaction of cysteine-modified proteins followed by their stable integration into lipid microdomains^5^. Thus we asked; i) whether RBC has the dynamic process of S-palmitoylation, and if it has, then how it regulates the defined localization and function of proteins and allow their precise tethering to membrane. To answer this query, we have used click chemistry and highly senstive chemico-proteomics based tool, namely ABE-coupled to LC-MS/MS. Click chemistry is based upon a simple chemical reaction that is triggered between palmitic acid alkyne analogues (17ODYA), which gets incorporated into exposed cysteines in membrane proteins, using cell’s own biochemical machinery and the azide functionalized dye (Oregon green), which helps in visualizing the cysteine medication by IFA^22^. Using this tool, we demonstrated for the first time ever, existence of a dynamic process of palmitoylation in RBC membrane, as clearly depicted by Oregon green stained RBCs (Figure-1(a)). Upon treatment with 2BP, RBCs demonstrated drastic reduction in green fluorescence, a typical signature of aborted palmitoylation process (Figure-1(a)). To identify the palmitoylated proteome and enrich the same from RBC ghost; we used ABE, a specialized cysteine-centric tool^23^ that allowed; i) *irreversible blocking of free thiols by alkylation*, ii) *breaking of thioester bonds in modified proteins by HA*, iii) *subsequent labelling of exposed thiols by sulfhydryl-reactive derivatives* (*biotin-HPDP*), followed by purification of modified proteins on streptavidin coated beads. Visualisation of silver stained gel represented appearance of several new protein bands in HA-treated RBC ghost, inferring to enriched pool of palmitoylated protein clusters. Upon blocking the PATs with 2BP, these bands got disappeared, depicting differential palmitoylation (Figure-1(b)). Further, LC-MS/MS-based analysis of global palmitoylome in RBC revealed 118 proteins, while 2BP-treated RBC ghost demonstrated 42 proteins (Figure-2(a)). Prediction of palmitoylation sites by CSS-Palm revealed more than one site per protein in ∼90% of the palmitoylome (Figure-2(b)). Delineating the functional classification revealed several clusters, pertaining to localization, plasma membrane, vesicle-mediated transport, transporter activity and secretory vesicles *etc* (Figure-2(c)). GO-enriched functional characterization of palmitoylome represented that, majority of proteins play important roles in maintaining cellular components; including, plasma membrane, cytoskeleton and cell surface, and also contribute in molecular processes, such as regulation, binding and catalytic activities (supplementary Figure-1). Interestingly, inhibiting PATs in RBC led to altered regulation of proteins belonging to transporters, eryptosis, cytoskeletal proteins, glycolysis and fatty acid metabolism (Figure-2(d)). Expression of proteins committed to anti-oxidant system, kinases, and vesiculation were non-detectable in 2BP-treated RBC ghosts (Figure-2(d)). These findings suggested that palmitoylation governs the functional classification of RBC membrane proteome and alteration in the same might lead to their deregulated spatio-temporal expression. Protein-protein interactome analysis exposed the majority of structural proteins and their possible interactions involved in crucial cellular processes (Supplementary Figure-2) and RBC-linked disease manifestation. Also, the structural palmitoylome (Figure-3(b)) was found to be linked to two clusters representing “haemolytic anaemia” and “Autosomal dominant inheritance” (Figure-2(e)). Additionally, we also found drastic depletion in DHHC17 expression in 2BP-treated RBC ghost (Supplementary Figure-3(b)). Thus, we assumed that the palmitoylation-dependent dynamic remodelling of structural proteins in RBC might be the foundation underlying it’s cytoskeletal flexibility. To explore this, we next analysed the profile of differential palmitoylome, which revealed that crucial structural proteins like; Spectrin α & β, Actin, Ankyrin, Band3, Protein4.2, Adducin1, Flotilin, CD44, CD151, Kell, the type II glycoprotein partner of Kell blood group system and others are dynamically palmitoylated. This finding suggested that RBC homeostasis and cytoskeletal deformability, might be governed by the dynamic palmitoylation status of its structural proteins. However, we also hypothesized that besides assigning flexibility, palmitoylaton might also play decisive role in making the RBC membrane susceptible to malaria parasite invasion. Our findings from global palmitoylomic profile strengthened this hypothesis, as Band3, ABCB6, Semaphorin7A and Complement Receptor1, the known molecular docking sites for parasite invasion^11, 12, 24–26^ are found to be significantly palmitoylated. Thus we assumed that, blocking palmitoylation might affect the parasite invasion via deregulation of host-derived factors. As assumed, abrogating the PATs in RBCs by 2BP treatment led to significant inhibition in invasion of *P. falciparum*, as with 100 μM of 2BP, parasitemia was diminished to 70%, suggesting a cumulative impact of abrupted palmitoylation on RBC membrane proteins, which are specifically involved in parasite invasion. Since vulnerability of RBC membrane is strongly influenced by the dynamicity of its structural proteins^27^, and majority of the structural proteins are found to be dynamically palmitoylated, we decided to look in-depth for a novel receptor proto-type for *P. falciparum* invasion in the same pool of structural palmitoylome, as shown in scheme (Figure-3(b)). Plausibly, our study unravelled that, among these proteins, Kell, a family of M13 Zinc metallo-endopeptidases demonstrated, i) negligible expression in RBC palmitoylome, ii) reduced expression in RBC ghost, and iii) complete disappearance from RBC membrane following 2BP-treatment, as depicted by LC-MS/MS, immunoblotting and IFA respectively. To emphasize, aborted palmitoylation might have led to dysregulated localization of Kell on RBC surface. Further, structural and functional analysis of Kell revealed some interesting guidance cues, such as; Kell, a 93 kDa glycoprotein is usually co-expressed with Kx as a part of Kell blood group system, and covalently linked to Kx via a crucial single disulphide bond between Cys**^72^**/Kell and Cys**^347^**/Kx^28^. Additionally, Kell has a unique exofacial antigenic determinant domain of 665 amino acids with a conserved HxxLH motif (histidine-glutamate-leucine-leucine-histidine), critical for its proteolytic activity. We hypothesized that Kell, being one of the dynamically palmitoylated skeletal proteins, with its unique structural attributes might play crucial role in *P. falciparum* invasion. To decipher the fundamental role of Kell in parasite interaction and invasion, we then treated RBC with Kell-specific antibody to ensure complete blockage of Kell activity, prior to *P. falciparum* invasion study. The result showed severe depletion in ring formation following exposure to Kell antibody for 6 hrs, which suggested that Kell is a potent receptor proto-type for parasite invasion (Figure-6(A-C)). Since stable expression of Kell on RBC surface is one of the pre-requisite conditions for parasite interaction, we revalidated its localized expression on RBC surface in sickle-cell RBCs from two SCA patients, the known malaria resistant RBC phenotype. As expected, these cells showed diminished expression of Kell on RBC surface with no change in Kx expression, with poor parasite invasion phenotype as compared to healthy control (Figure-6(D)). To further find a possible correlation between the altered palmitoylation status with distorted expression of Kell on sickle-cell RBCs, we performed ABE to extract palmitoylated proteins from sickle-cell RBC ghost lysate and analyzed the Kell expression by immunoblotting. As assumed the palmitoylated Kell was drastically reduced in sickle-cell RBCs (Figure-5(d)). Since Kell_null_ individuals have no clinical manifestations and absence of Kell has no impact on structural and functional significance of RBC^29^, we propose Kell as potent host-specific candidate for developing novel anti-malarial chemotherapeutics in future. Overall, this study reports three novel findings (Figure-6), namely; i) regulated expression of RBC membrane proteins is strictly governed by its dynamic palmitoylation status, ii) palmitoylation-dependent localized expression of Kell on membrane introduces it as a potent parasite-specific host receptor, and iii) altered palmitoylation-mediated dysregulated localized expression of Kell on sickle-cell RBCs might be one of the reasons underlying their resistance to malaria.

**Figure 6.**
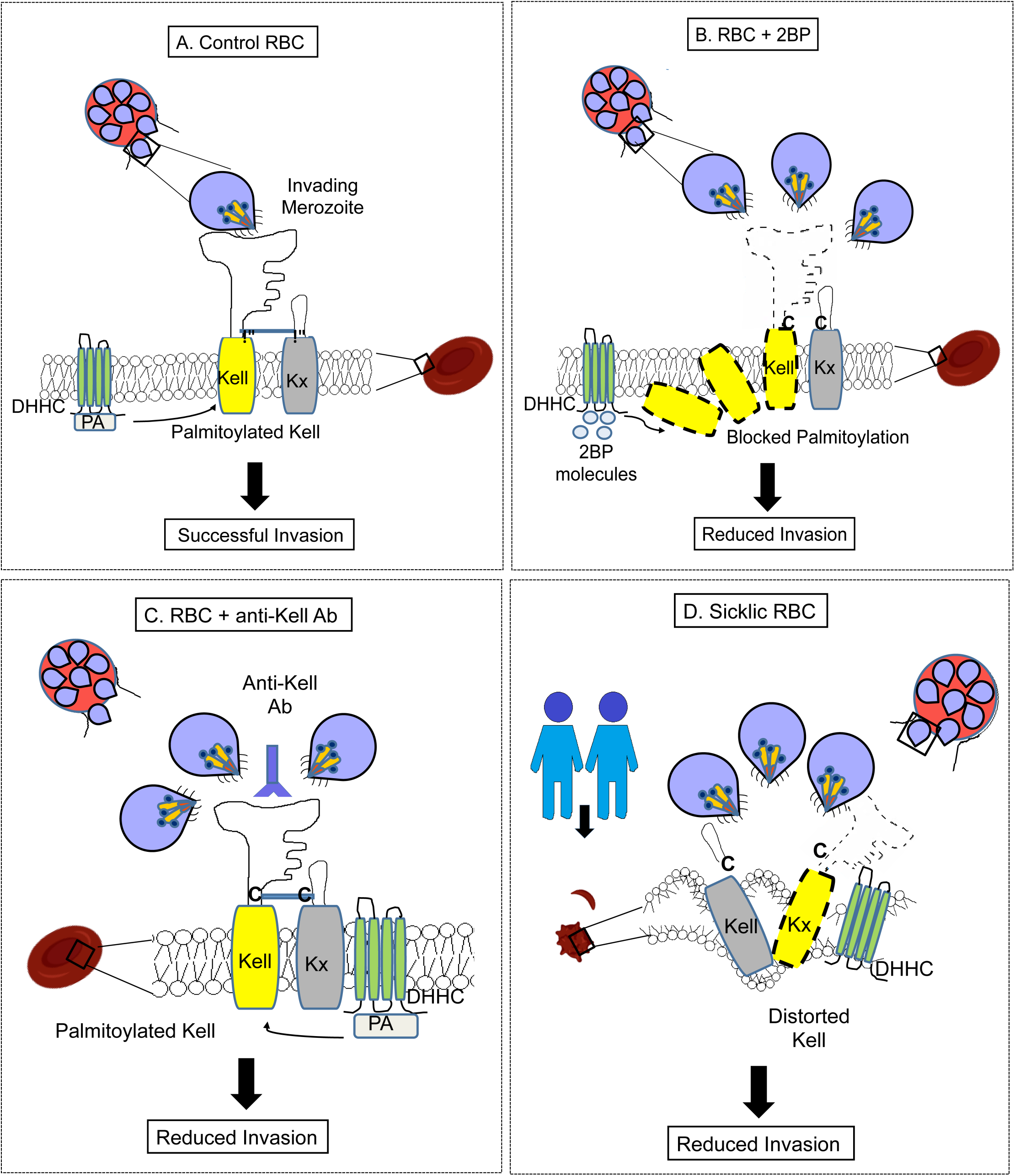
Scheme represents Kell-Kx organization in RBC membrane under different conditions (A-D), and depicts the role of Kell in *P.falciparum* invasion.

## Methods

### Parasite culture

Frozen stock of *P. falciparum* 3D7 line was thawed and cultured in human O+ erythrocytes at 2% haematocrit under mixed gas condition (5% CO_2_, 5% O_2_, 95% N_2_) at 37°C as per the standard protocol previously reported from our laboratory^30^. Culture was grown in medium containing RPMI (RPMI 1640, Invitrogen) supplemented with 2g/L NaHCO_3_ (Sigma), 5g/L Albumax (Invitrogen, Waltham, MA, USA), 50mg/L Hypoxanthine (Sigma), 10µg/mL gentamicin sulfate (Invitrogen).

### Click Chemistry-based RBC palmitoylome labelling

The palmitoylation status of RBCs was examined by labelling the cells with Click Tag^TM^ Palmitic Acid Alkyne, a 16-carbon saturated fatty acid group as described in our previously published article^15, 31^. Briefly, palmitic acid (Alk-C16, Cayman Chemicals) was dissolved in DMSO to achieve final stock concentration of 20 mM. Cells were washed twice with ice-cold PBS, fixed with chilled methanol for 5 min, and further permeabilized using 0.01% Triton X-100 (Sigma) in PBS at RT for 5 min. These processed bacterial samples were subjected to a click labelling reaction in 100 µL of dye mix containing 0.1 mM azide dye (Oregon Green 488-azide, Thermo Fisher Scientific, Waltham, MA, USA), 1 mM Tris-(2-carboxyethyl)-phosphine hydrochloride (TCEP-Sigma), and 1 mM CuSO4 (Sigma) in PBS for 1 h. After incubation, cells were pelleted down and washed twice by 1X PBS. For microscopy, a smear was made which was mounted in antifade mounting solution. Images were acquired by fluorescence microscope (Zeiss, Oberkochen, Germany).

### Acyl biotin exchange (ABE) method to purify RBC Membrane palmitoylome

Purity of RBC samples is a prerequisite for their unbiased proteome analysis. Packed RBCs were procured from Blood Bank and were washed thrice using incomplete RPMI 1640 medium to remove any remnant debris and cell types. 1mL packed RBCs were washed twice using 1X iRPMI (incomplete RPMI) and resuspended in it to maintain 50% haematocrit. For 2BP treatment, cells were treated with 100 µM of 2-BP (Sigma) and incubated overnight at 37^0^C. RBCs were then washed twice in 1X PBS, resuspended in ice-cold 0.1X PBS hypotonic solution, followed by repeated freeze-thaw cycles. These lysis steps were repeated until white RBC ghosts were obtained. Extraction of membrane protein from RBC ghosts was carried out by solubilizing 1 volume of packed RBC ghosts with 4 volumes of 50mM HEPES (pH 7.4), 1% Triton-X 100, 1% NP-40, protease inhibitor cocktail and incubating the mixture at 4^0^C for 16 hrs with 10mM NEM. Detergent sensitive and detergent resistant fractions were separated by centrifugation at 13,000 rpm for 30min at 4^0^C. The detergent-sensitive fraction was processed for ABE using the modified method previously reported^15, 32^. Samples were run on 10% SDS gel, and visualised using silver staining. For immunoblotting, proteins were transferred to nitrocellulose membrane and blotting was done using anti-Kell antibody. For ABE of sickle-cell RBCs, blood was obtained from a 10-year-old male and a 9-year-old female patient, and was processed according to the below mentioned protocol.

### Sickle cell patients’ blood collection and RBC separation

Sickle cell anemia patients’ peripheral blood samples were collected in EDTA tubes following Institutional human ethical approval (IHEC) and informed patient consent. The blood samples were diluted with phosphate buffered saline (PBS) and processed for cell separation using Ficoll-Paque™ (GE healthcare) based density gradient centrifugation method. The blood cells were separated based on their respective densities in the tube: platelets at the top, peripheral blood mono-nuclear cells (PBMCs) at the middle and red blood cells (RBCs) at the bottom of the tube. First two cell fractions (platelets and PBMCs) were removed and RBCs were collected and washed with 1X PBS (Phosphate Buffer Saline) and used for further analysis.

### In-solution digestion

Briefly, eluted proteins were vacuum dried and resuspended in 1 M guanidium hydrochloride (GmCl). To remove the GmCl, ethanol precipitation was carried out with 9 volumes of chilled 100% ethanol for 60 min at –20°C. Eluted Samples (n=4) were further centrifuged at 13000xg for 20 min at 4^0^C and then supernatant was carefully removed. Pellet was washed with chilled 90% ethanol (kept at 20^0^ degree) by centrifugation at 13000xg for 10 min at 4^0^C. Finally protein pellets were dried in SpeedVac concentrator (Thermo Scientific) and resuspended in 50 mM TEABC buffer. Samples were reduced with 10 mM dithiothreitol (DTT) for 20 min at 60^0^C and alkylated for 10 min at room temperature in dark, using 20 mM iodoacetamide (IAA), respectively. The protein was digested with trypsin (1:20, W/W) (Promega, Madison, WI) at 37°C for 16 hours. The reaction was stopped by adding 0.1% formic acid, followed by desalting using SCX (Strong Cation Exchange) StageTips after which the eluted peptides were lyophilized at room temperature and stored at −20^0^C until LC-MS/MS analysis.

### Liquid Chromatography Mass Spectrometry (LC-MS/MS analysis)

LC-MS/MS based identification of RBC palmitoylome was done based on an established protocol^33^. The peptides were analyzed on Orbitrap Fusion Tribrid mass spectrometer (Thermo Fischer Scientific, Bremen, Germany) connected to Easy-nLC-1200 nanoflow liquid chromatography system (Thermo Scientific). The peptides were reconstituted in 0.1% formic acid and loaded onto a trap column (Thermo Scientific, 75 µm X 2 cm, nanoViper, 3 µm, 100 Å filled with C18 material, at a flow rate of 4 µL/min. Peptides were separated using a 15 cm analytical column (EASY-SPRAY RSLC C18 2 µm 15X50 µm; Thermo Scientific) at a flow rate of 300 nl/min. The solvent gradients were set as the linear gradient of 5-35% solvent B (80% acetonitrile in 0.1% formic acid) over 90 min and total run time of 120 min. MS analysis was carried out at a scan range of 400-1600 m/z mass range (120,000 mass resolution at 200 m/z) in a data-dependent mode using an Orbitrap mass analyzer. The maximum injection time was 10 ms. For MS/MS analysis, data was acquired at top speed mode with 3s cycles and subjected to higher collision energy dissociation (HCD) with 32% normalized collision energy. MS/MS scans were carried out at a range of 100-1600 m/z using Orbitrap mass analyzer at a resolution of 30,000 at 200 m/z. Maximum injection time was 200ms.

### Data Processing Protocol and Data analysis

Mass spectrometry-derived data was searched against the reference protein database *Human_RefSeq92* (consisting of 81,064 protein entries along with common contaminants) from NCBI. The mass spectrometry data was analyzed with SEQUEST HT and Mascot (version 2.5.1; Matrix Science, London, United Kingdom) search algorithms in Proteome Discoverer software suite, version 2.2 (PD 2.2) (Thermo Fischer Scientific, Bremen, Germany). The search parameters used were; a) trypsin as the proteolytic enzyme (with up to one missed cleavage); b) fragment mass error tolerance of 0.05 Da); c) peptide mass error tolerance of 10 ppm; d) oxidation of methionine as a variable modification; e) carbamidomethylation of cysteine as a fixed modification. A false discovery rate (FDR) of 1% was applied while identifying the peptide-spectrum matches (PSMs).

### Annotation of GO and Functional bioprocess analysis

To explore the function of enriched proteins, we performed GO analysis using g:Profiler. The g:GOSt tool in the g:Profiler suite uses a Fisher’s one-tailed test to measure the statistical significance of enrichment for any given GO term. Multiple testing is corrected using the built-in previously described g:SCS^34^ and STRAP^35^ methods, to classify proteins based on their Information about biological processes (BP), molecular functions (MF), and cellular components (CC). The phenotypes are organized into structured vocabularies which are automatically obtained from Gene Ontology (GO), Human Phenotype Ontology (HPO), Reactome, and KEGG molecular pathways.

### Protein Protein Interaction (PPI) network analysis

Prediction of protein interactome pathways was carried out using STRING (Search Tool for the Retrieval of Interacting Genes/Proteins) (http://string-db.org/)^36^. The parameters used for the STRING analysis included interaction sources from databases and literature, analyses of co-expressed genes and also from computational predictions to identify interacting partners as associated networks.

### Computational prediction of palmitoylated proteins and motif

We used the mass-spectrometry proteomics dataset to identify palmitoylome in both samples. The palmitoylome was characterized by predicting palmitoylation sites by applying CSS-Palm 4.0^37^ based clustering and scoring strategy (http://csspalm.biocuckoo.org/).

### Immunoblotting

To check the expression of various RBC membrane proteins, the detergent sensitive fraction of RBC ghost was electrophoresed on 10% SDS-gel, and transferred to 0.45-um nitrocellulose membrane (Bio-Rad) in a semi-dry transfer apparatus (Bio-Rad). This protocol was modified based on the earlier published articles from our laboratory^15, 38^. Non-specific sites were blocked by incubating the membrane with 5% fat-free skimmed milk powder (prepared in 1X PBST), for 1hr at room temperature. After incubation, blot was washed thrice using 1X PBS and was then incubated with specific primary antibodies. For detection of Kell, Kx, MPP1, Protein4.1 and DHHC17; primary antibodies used were goat-anti-Kell (Invitrogen, 1:2000 dilution), rabbit-anti-Kx (Invitrogen, 1:2000 dilution), rabbit-anti-MPP1 (Bethyl Laboratories, 1:500 dilution), rabbit-anti-Protein4.1 (Invitrogen, 1:1000 dilution) and rabbit-anti-DHHC17 (Invitrogen, 1:1000 dilution) respectively for overnight at 4^0^C. Following washing, the nitrocellulose membrane was incubated with respective HRP-conjugated secondary antibodies. Immunoblots were analysed using enhanced chemiluminescent system (ECL).

### Immunofluorescence assay (IFA)

To perform IFA in both treated and untreated RBCs, we have used standardized protocol based an earlier report from our laboratory^38^. To prepare RBC samples for IFA, 10uL packed RBCs (from healthy human donor or SCA patient) were washed twice with 100uL of 1X iRPMI, supernatant was aspirated off and the cells were resuspended in 100uL iRPMI. To check the expression of Kell and Kx on surface of RBCs in the presence and absence of 2-BP, the cells were treated with 100 µM 2-BP and incubated overnight at 37^0^C, while the control and sickle-cells (malaria resistant phenotype) were left untreated. After incubation, cells were washed twice in 1X PBS, fixed in 0.25% glutaraldehyde at 4°C for 15min. Following incubation, cells were washed with 1X PBS and permeabilized with 0.01% Triton X-100 at RT for 5 minutes. Cells were washed again with 1XPBS and treated with 200uL of goat-anti-Kell and rabbit-anti-Kx antibodies for 2hr at room temperature. Washing was followed by incubation with anti-goat-Alexa-Fluor488 and anti-rabbit-Alexa-Flour488-conjugated secondary antibodies for detection of Kell and Kx respectively for 1hr at room temperature in dark. Incubation was followed by washing with 1X PBS and the cells were suspended in 100uL of 1X PBS. For microscopic analysis a smear was drawn, mounted in anti-fade mounting solution and was visualised using confocal microscope (Nikon).

### Invasion Assay

To study parasite invasion, we have used an already established protocol from our laboratory^39^, in which *P. falciparum* 3D7 strain was Percoll-purified to enrich the 44-46 hour schizont-stage parasites, with 95% purity and cultured in 96-well plate in a total volume of 50uL. Haematocrit and initial parasitemia were adjusted to 2% and 1% respectively. To study the role of Kell in parasite invasion, in-house generated anti-Kell IgG was added to the culture wells at different concentration (0.02, 0.04, 0.06 mg/mL). To study the role of 2BP on parasite invasion, uninfected RBCs were pre-incubated with 2BP for 12 hours at different concentrations and were then used to maintain 2% haematocrit. Untreated control cells were used to validate the normal invasion ability of the parasite, while sickle-cells were used to study the malaria resistant phenotype. After 20hr, newly invaded ring-stage parasites were counted, by making a smear and staining with Giemsa (Sigma).

### Statistical analysis

The data for all the assays are expressed as the mean ± standard deviation (SD) of three independent experiments done in triplicates. Student’s *t*-test (one-tailed) was performed to calculate the p values, where p < 0.05 was taken as significant.

### Data Availability

All data are available from the authors upon reasonable request.

## Acknowledgements

We are thankful to Advanced Instrumentation and Research Facility (AIRF), JNU, New Delhi for confocal microscopy, Yenepoya University for access to instrumentation and Central Instrumentation Facility (CIF) of Special Centre for Molecular Medicine, JNU for other instruments and facilities. This work has been funded by Innovative Young Biotechnologist Award (IYBA) from the Department of Biotechnology, Ministry of Science and Technology, Government of India (DBT); National Institute of Health (Grant No. U19AI089676-09), USA and DST-EMR (SERB, File No. EMR/2016/005644) grant from Department of Science & Technology. S.S. is a recipient of the IYBA Award from DBT. SP is greatful to support from Shiv Nadar Foundation. GK is a recipient of Junior Research Fellowship from Council of Scientific and Industrial Research, Government of India. PY is a recipient of Senior Research Fellowship from University Grants Commission, Government of India. Rex is a recipient of Senior Research Fellowship from Indian Council of Medical Research, Government of India. The funders had no role in study design, data collection and analysis, decision to publish, or preparation of the manuscript.

## Author Information

These authors contributed equally: Soumya Pati, Preeti Yadav, Geeta Kumari

## Contributions

S.S. conceived the idea and designed the experiments, wrote and edited the manuscript. S.P and P.Y executed the experiments, performed the data interpretation, analyzed the experimental and palmitoylome data, and wrote the manuscript. R.D. and T.P. performed Mass Spectrometry, R.D. performed bioinformatics analysis. G.K. & S.G. conducted microscopy experiments. S.G. (*IGIB*) and S.R. provided SCA samples and related methodologies.

## Corresponding author

Shailja Singh.

## Abbreviations

RBC: Red Blood Cell
LC-MS/MS: Liquid Chromatography with Tandem Mass Spectrometry
IFA: Immunofluorescence Assay
PTM: Post-translational modification
HA: Hydroxylamine
MFI: Mean Fluorescence Intensity
17-ODYA: 17-Octadecynoic acid
HPDP: (N-[6-(biotinamido)hexyl]-3’-(2’-pyridyldithio)propionamide)
SCA: Sickle Cell Anaemia
GO: Gene Ontology

