## Supplemental Information for "Role of RBC membrane protein palmitoylation in regulation of molecular topology and susceptibility to *Plasmodium falciparum* invasion"

**This PDF file includes:**

Figures S1 to S4
Tables S1 to S2
Legends for Figures S1 to S4
Legends for Tables S1 to S2

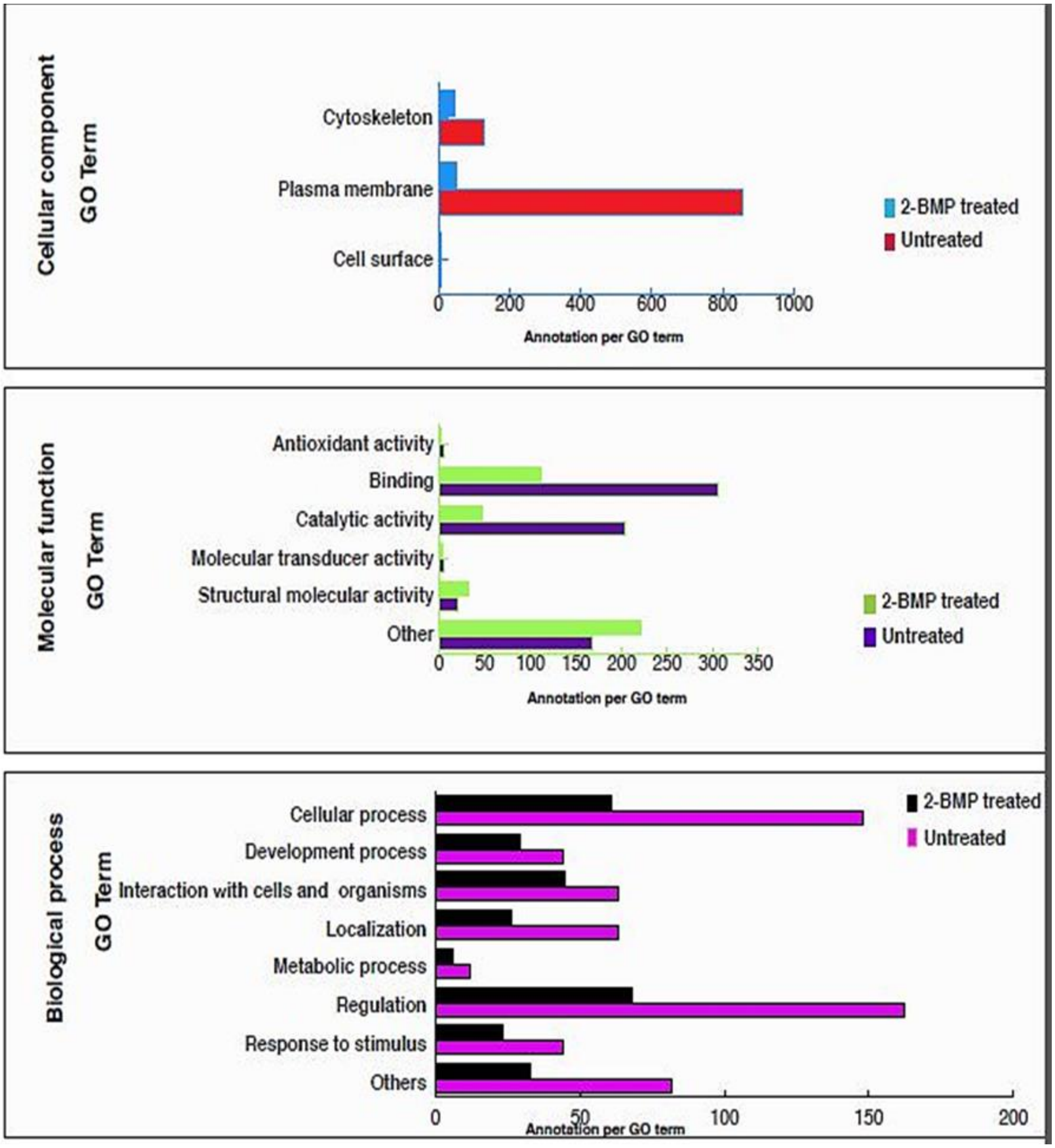


Fig. S1. Gene Ontology analysis of predicted palmitoylated proteins of RBC membrane highlighting the cellular component, molecular function and biological process relevant to RBC biology.

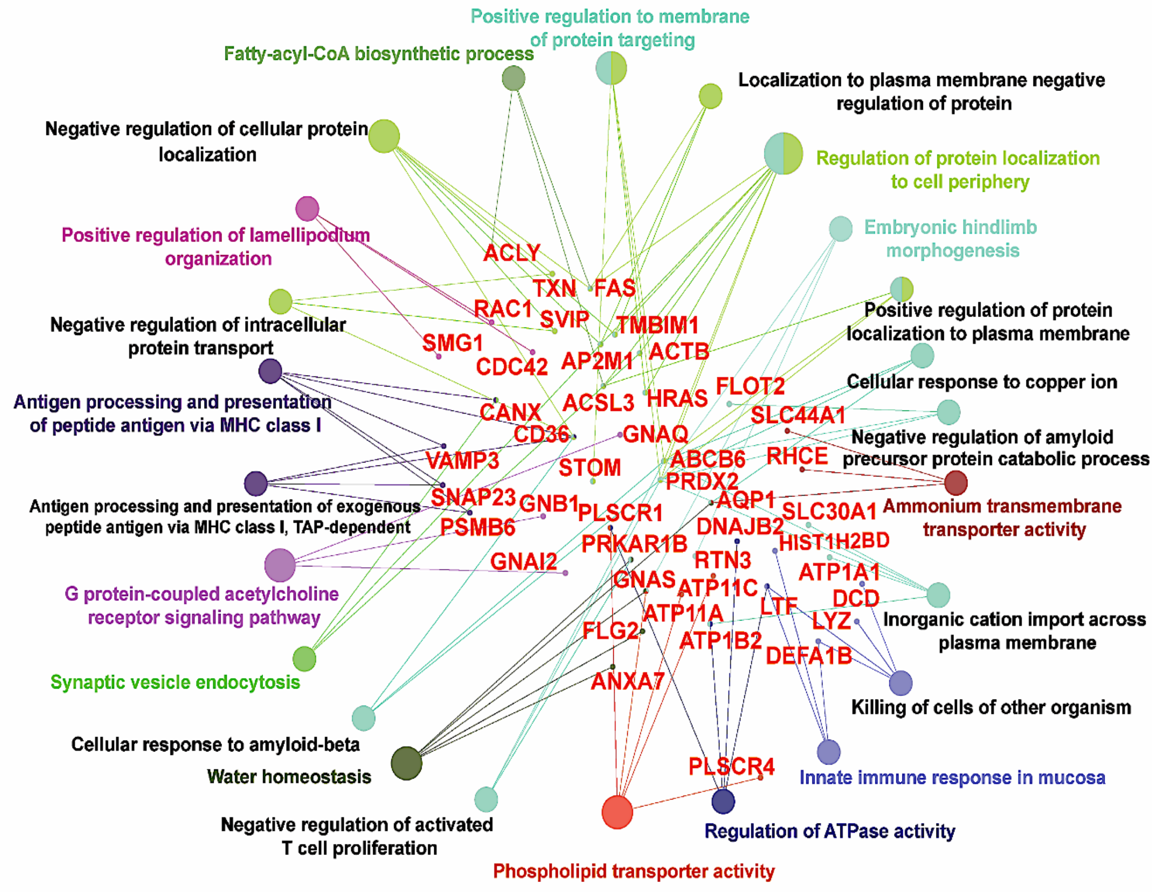


Fig. S2. Diagram depicting the functional classification and interaction of RBC ghost palmitoylome.

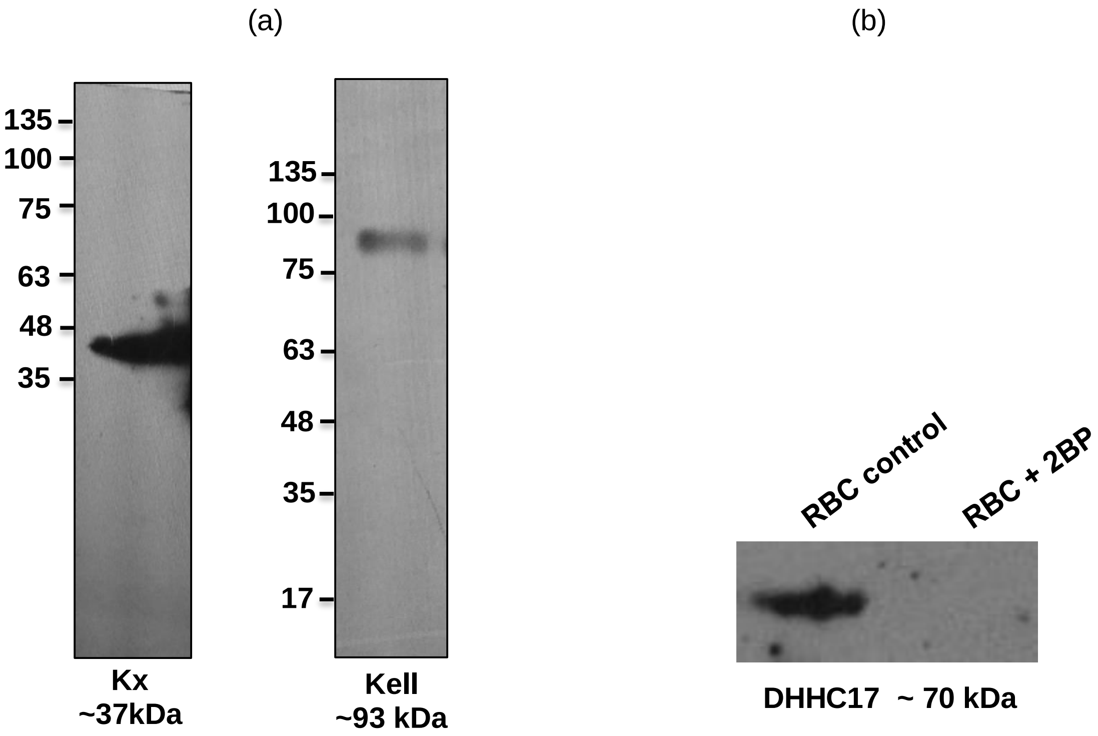


Fig. S3. (a) Immunoblots depicting the expression of Kell and Kx in RBC ghost lysate using goat-anti-Kell and rabbit-anti-Kx IgGs respectively. (b) Immunoblot showing the expression of DHHC17 in control untreated and 2BP-treated RBC ghost lysate.

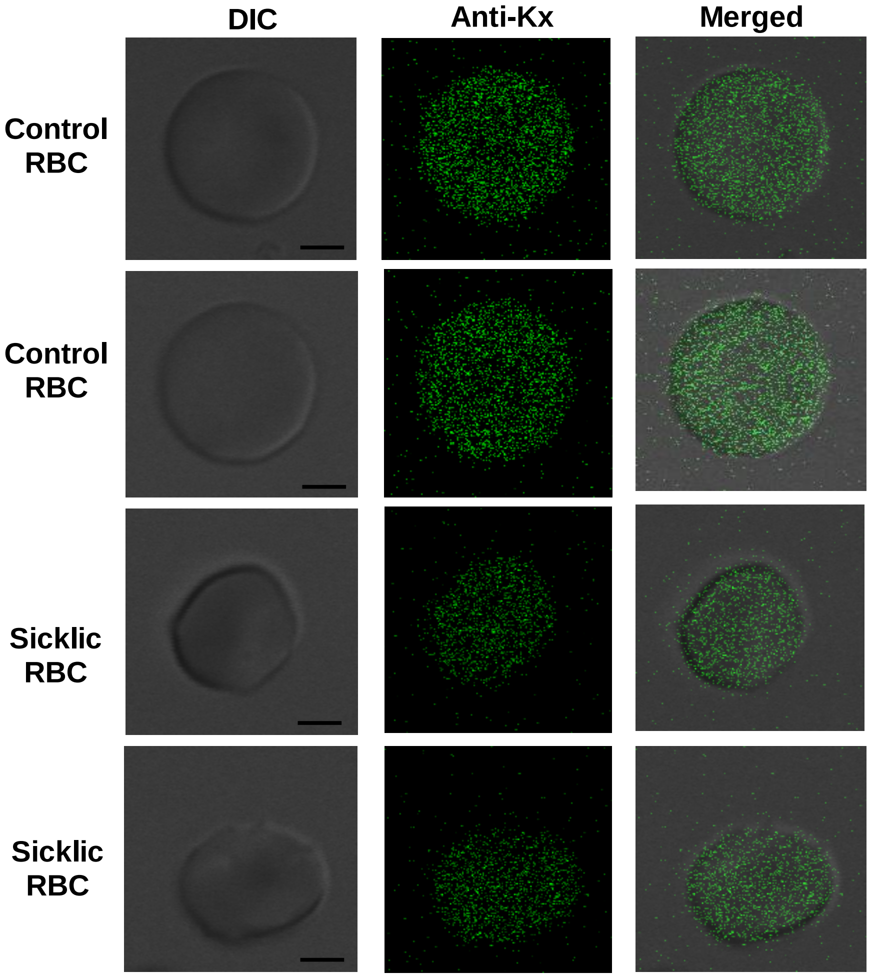


Fig. S4. Confocal images showing localisation of Kx on surface of sickle-cell RBCs, in comparison to normal healthy RBCs.

| Accession No. | Protein Name | Coverage (%) | Score | Unique peptides | PSM |
|---|---|---|---|---|---|
| NP_001341087.1 | Acyl-CoA synthetase long chain family member 3 | 2 | 4.132 | 1 | 4 |
| XP_011522750.1 | adaptor related protein complex 2 | 1 | 2.374 | 1 | 3 |
| NP_004025.1 | Annexin-A7 | 2 | 2.966 | 1 | 9 |
| NP_001316801.1 | Aquaporin-1 | 6 | 8.216 | 1 | 4 |
| NP_001231367.1 | Arginase-1 | 5 | 5.873 | 2 | 14 |
| XP_011515270.2 | Argonaute | 5 | 16.402 | 4 | 21 |
| XP_016884928.1 | ATPase phospholipid transporting 11C | 2 | 5.255 | 3 | 12 |
| XP_011542295.1 | calcium/calmodulin dependent serine protein kinase | 2 | 5.721 | 1 | 4 |
| NP_001675.3 | Calcium transporting ATPase | 7 | 46.885 | 4 | 106 |
| XP_024301993.1 | Calnexin | 1 | 2.918 | 1 | 2 |
| NP_001185797.1 | Calpain-1-catalytic subunit | 1 | 1.672 | 1 | 3 |
| NP_001158230.1 | cAMP dependent protein kinase type-1 beta | 3 | 4.241 | 1 | 2 |
| NP_056278.1 | Cannabinoid receptor interacting protein 1 | 15 | 9.355 | 2 | 18 |
| NP_000058.1 | Carbonic anhydrase -2 | 3 | 2.331 | 1 | 2 |
| NP_001766.2 | CD38 | 7 | 6.237 | 2 | 14 |
| NP_000602.1 | CD59 antigen | 6 | 1.676 | 1 | 3 |
| XP_005257725.1 | Charged multivesicular body protein-6 | 5 | 4.17 | 1 | 2 |
| XP_024309055.1 | Complement C3b/C4b receptor-1 | 2 | 3.824 | 1 | 4 |
| NP_000177.2 | Complement factor H precursor | 1 | 6.053 | 1 | 6 |
| NP_001329.1 | Coxsackie virus and adenovirus receptor precursor | 4 | 1.834 | 1 | 4 |
| NP_149124.3 | Cyclic nucleotide phosphodiesterase precursor | 8 | 16.569 | 3 | 18 |
| NP_061933.1 | Cyesteine rich C-terminal protein-1 | 13 | 3.675 | 1 | 3 |
| NP_079119.3 | Cytochrome-b reductase 1 | 3 | 3.86 | 1 | 8 |
| NP_085056.2 | Cytochrome-B-5-type-B precursor | 8 | 2.545 | 1 | 2 |
| NP_001017922.1 | Erythroblast membrane associated protein | 3 | 5.212 | 2 | 11 |
| NP_001071643.1 | Equilibrated nucleoside transporter 1 | 7 | 11.329 | 3 | 17 |
| NP_001435.1 | Fatty acid binding protein 5 | 15 | 6.422 | 2 | 11 |
| NP_001230106.1 | Fructose biphosphate aldolase | 10 | 15.552 | 4 | 41 |
| NP_001344607.1 | Galectin-3 | 3 | 1.806 | 1 | 3 |

| | | | | | |
|---|---|---|---|---|---|
| NP_033665.1 | Galectin-9 | 4 | 5.412 | 1 | 6 |
| NP_001034937.1 | Glutathione peroxidase | 3 | 1.981 | 1 | 2 |
| NP_002073.2 | GPCR kinase 6 | 3 | 6.727 | 2 | 30 |
| NP_001269468.1 | Guanine nucleotide binding protein | 6 | 9.911 | 1 | 26 |
| NP_056272.2 | Human DHHC5 | 3 | 7.234 | 2 | 10 |
| NP_055400.1 | Human ferroportin | 7 | 22.637 | 4 | 62 |
| NP_001535.1 | ICAM -4 | 14 | 19.895 | 3 | 23 |
| NP_002334.2 | Lactotransferrin precursor | 1 | 2.806 | 1 | 4 |
| XP_005246300.1 | LanC lantibiotic synthetase component-C-like-1 | 2 | 3.141 | 1 | 2 |
| XP_006721660.1 | Mannose-P-dolichol utilization defect-1 | 4 | 2.99 | 1 | 2 |
| NP_006311.2 | Membrane-associated progesterone receptor-component-2 | 8 | 2.832 | 1 | 2 |
| NP_054752.3 | Methyltransferase like protein 7A precursor | 5 | 5.312 | 1 | 8 |
| **NP_005836.2** | Multidrug resistance associated protein 4 | 1 | 4.572 | 2 | 12 |
| NP_005800.3 | Peroxiredoxin-2 | 9 | 6.335 | 2 | 24 |
| NP_060895.1 | Phosphatidylinositol-4-kinase-type-2-alpha | 7 | 12.867 | 3 | 56 |
| NP_000012.1 | Presenilin | 2 | 3.774 | 1 | 4 |
| NP_001310245.1 | Protein glutamine gamma-glutamyltransferase-2 | 2 | 2.326 | 1 | 2 |
| NP_001159477.1 | Protein 4.1 | 26 | 93.742 | 2 | 236 |
| NP_002659.1 | Proteolipid protein 2 | 10 | 6.983 | 1 | 4 |
| NP_002789.1 | Proteosome subunit beta | 5 | 3.707 | 1 | 2 |
| XP_005268509.2 | Protocadherin-1 | 1 | 1.92 | 1 | 3 |
| NP_001238965.1 | RAB5B | 6 | 4.202 | 1 | 13 |
| NP_001238968.1 | RAB5C | 5 | 5.247 | 1 | 10 |
| NP_055814.1 | RAB21 | 11 | 7.01 | 2 | 15 |
| NP_001306000.1 | SAC1 like phosphatidylinositide phosphatase | 4 | 6.035 | 3 | 11 |
| XP_016862117.1 | Scramblase 1 | 9 | 13.835 | 3 | 60 |
| NP_003603.1 | Semaphorin-7A precursor | 1 | 1.794 | 1 | 2 |
| NP_003013.1 | SH3-domain binding-glutamic-acid rich like protein | 11 | 2.021 | 1 | 6 |
| NP_000692.2 | Sodium/pottassium transporting ATPase subunit | 1 | 4.701 | 2 | 14 |
| NP_001159968.1 | Solute carrier family 16 | 2 | 2.314 | 1 | 20 |

| | | | | | |
|---|---|---|---|---|---|
| NP_001185739.1 | Solute carrier family 43 member 1 | 2 | 3.568 | 1 | 1 |
| XP_006717090.1 | Solute carrier family 44 member 1 | 2 | 6.198 | 1 | 9 |
| XP_001614990.1 | Suppressor of RAS-1 3.9 putative | 3 | 1.653 | 1 | 8 |
| NP_110382.3 | Theoredoxin related transmembrane protein 1 | 12 | 12.445 | 3 | 34 |
| NP_003320.2 | Thioredoxin | 9 | 3.263 | 1 | 15 |
| NP_001308356.1 | Transmembrane BAX inhibitor motif contatining 1, isoform | 5 | 7.019 | 1 | 2 |
| XP_024306385.1 | Transmembrane channel like 8 | 1 | 1.913 | 1 | 2 |
| NP_006125.2 | Transmembrane protein 50B | 7 | 3.33 | 1 | 21 |
| NP_955369.1 | Transmembrane protein 179 B | 4 | 1.844 | 1 | 4 |
| NP_115501.2 | Transmembrane protein 222 | 36 | 39.625 | 4 | 65 |
| NP_001159588.1 | Tropomodulin 1 | 4 | 11.509 | 1 | 21 |
| NP_001317273.1 | Tropomyosin 1- alpha | 23 | 25.871 | 6 | 77 |
| XP_016858330.1 | Vesicle associated membrane protein 3 | 16 | 4.719 | 2 | 12 |
| NP_955376.1 | Vesicle transport protein SFT2B | 6 | 4.585 | 1 | 13 |
| NP_001335099.1 | WW domain binding protein 2 | 5 | 7.347 | 1 | 10 |
| NP_001176.1 | Zinc-alpha-2-glycoprotein precursor | 3 | 3.503 | 1 | 2 |
| NP_067017.2 | Zinc transporter 1 | 5 | 8.564 | 2 | 6 |

Table S1. List of palmitoylated proteins unique to control RBC ghost fraction that were completely absent from 2BP treated RBC ghost

| Accession No. | Protein Name | Control RBCs | | | | 2BP treated RBCs | | | |
|---|---|---|---|---|---|---|---|---|---|
| | | Coverage (%) | Score | Unique peptides | PSM | Coverage (%) | Score | Unique peptides | PSM |
| NP_001092.1 | Actin | 25 | 69.95 | 7 | 109 | 7 | 14.14 | 2 | 45 |
| NP_054908.2 | Alpha - adducin | 9 | 20.3 | 7 | 36 | 2 | 2.431 | 1 | 4 |
| XP_011542792.1 | Ankyrin -1 | 23 | 216.1 | 34 | 358 | 11 | 24.91 | 5 | 181 |
| NP_000333.1 | Band -3 | 28 | 166.2 | 19 | 520 | 8 | 22.85 | 7 | 37 |
| NP_004090.4 | Band -7 | 42 | 96.97 | 10 | 361 | 15 | 14.57 | 4 | 147 |
| NP_005572.2 | Basal cell adhesion molecule precursor | 19 | 36.47 | 8 | 124 | 3 | 9.509 | 2 | 70 |
| NP_001171983.1 | Beta - adducin | 13 | 36.51 | 8 | 159 | 8 | 14.42 | 4 | 64 |
| NP_059118.2 | Calmodulin – like protein 5 | 15 | 4.439 | 2 | 8 | 9 | 4.873 | 1 | 12 |
| XP_011543527.1 | Calpain -5 | 8 | 26.29 | 6 | 89 | 1 | 3.904 | 1 | 25 |
| NP_001122301.1 | Carbonic anhydrase -1 | 14 | 17.68 | 3 | 11 | 3 | 2.638 | 1 | 6 |
| NP_001107607.1 | Dematin | 24 | 26.13 | 7 | 78 | 2 | 3.251 | 1 | 12 |
| NP_004406.2 | Desmoplakin | 2 | 19.1 | 6 | 40 | 1 | 6.353 | 2 | 5 |
| NP_068778.1 | E3 ubiquitin protein ligase | 1 | 2.388 | 1 | 2 | 20 | 4.79 | 1 | 1 |
| NP_006507.2 | Facilitated glucose transporter | 7 | 36.45 | 4 | 44 | 2 | 2.461 | 1 | 12 |
| XP_011538066.1 | Fas cell-surface death receptor | 2 | 3.116 | 1 | 39 | 2 | 2.277 | 1 | 24 |
| NP_005794.1 | Flotillin -1 | 49 | 115.2 | 22 | 446 | 7 | 10.3 | 3 | 125 |
| XP_024306434.1 | Flotillin -2 | 32 | 79.97 | 17 | 270 | 3 | 4.032 | 2 | 63 |
| NP_001276674.1 | GAPDH | 17 | 19.57 | 5 | 91 | 2 | 1.811 | 1 | 26 |
| XP_016883301.1 | GNAS complex locus | 4 | 13.15 | 3 | 78 | 2 | 6.385 | 2 | 51 |
| NP_001123914.1 | HRAS | 10 | 5.852 | 2 | 9 | 4 | 1.704 | 1 | 10 |
| NP_001017963.2 | HSP 90 | 2 | 5.848 | 1 | 21 | 2 | 9.836 | 1 | 84 |
| XP_005253288.1 | Indian Blood group | 9 | 39.79 | 6 | 136 | 1 | 1.892 | 1 | 7 |

| | | | | | | | | | |
|---|---|---|---|---|---|---|---|---|---|
| NP_058642.1 | Junctional adhesion molecule precursor | 4 | 3.038 | 1 | 4 | 3 | 1.785 | 1 | 2 |
| NP_000411.1 | Kell | 9 | 23.14 | 6 | 109 | 2 | 4.035 | 1 | 17 |
| NP_066569.1 | Kx | 6 | 13.32 | 3 | 33 | 4 | 4.637 | 2 | 30 |
| NP_005680.1 | Langereis Blood group | 16 | 64.79 | 10 | 116 | 1 | 11.06 | 1 | 55 |
| NP_001185610.1 | Leptin-receptor gene related protein | 9 | 3.575 | 1 | 14 | 9 | 5.64 | 1 | 15 |
| NP_002427.1 | MPP-1 | 47 | 154.9 | 16 | 466 | 10 | 22.3 | 4 | 79 |
| NP_000110.2 | Protein 4.2 | 22 | 128.2 | 16 | 437 | 4 | 6.765 | 2 | 9 |
| XP_016857449.1 | Rap1A | 9 | 6.968 | 2 | 30 | 4 | 1.614 | 1 | 10 |
| NP_001034579.1 | Raph blood group | 3 | 3.184 | 1 | 16 | 3 | 1.95 | 1 | 5 |
| XP_016865333.1 | Receptor accessory protein 5 | 3 | 2.121 | 1 | 6 | 3 | 1.781 | 1 | 9 |
| XP_011540191.2 | Rh blood group CcEe antigen | 4 | 5.056 | 2 | 16 | 2 | 1.772 | 1 | 12 |
| NP_001121776.1 | Scramblase - 4 | 9 | 13.42 | 3 | 64 | 2 | 2.062 | 1 | 6 |
| NP_001157196.1 | Small integral membrane protein 1 | 36 | 38.77 | 2 | 53 | 1 | 5.212 | 1 | 12 |
| NP_001156467.1 | Small integral membrane protein - 5 | 13 | 13.27 | 1 | 14 | 13 | 11.18 | 1 | 14 |
| NP_001265135.1 | Solute carrier family 43 member 3 | 3 | 4.983 | 2 | 10 | 2 | 2.408 | 1 | 7 |
| NP_003117.2 | Spectrin - alpha | 13 | 119.3 | 26 | 444 | 3 | 22.25 | 8 | 114 |
| NP_001020029.1 | Spectrin - beta | 8 | 83.62 | 17 | 202 | 1 | 8.278 | 2 | 17 |
| XP_016878181.1 | Synaptosome associated protein 23 | 33 | 22.05 | 5 | 33 | 5 | 5.331 | 1 | 16 |
| NP_001036816.1 | Tropo-myosin – alpha3 | 20 | 24.4 | 5 | 49 | 4 | 3.267 | 1 | 4 |
| NP_001139508.2 | Urea transporter | 9 | 14.75 | 4 | 59 | 5 | 7.806 | 2 | 34 |

Table S2. List of palmitoylated proteins common to both control and 2BP-treated ghosts, identified by LC-MS/MS upon ABE.
